# Neuronal processing of facial expressions in the primate superior colliculus

**DOI:** 10.64898/2026.09.10.750132

**Authors:** Gongchen Yu, Leor Katz, Richard J. Krauzlis

## Abstract

Fast processing of emotionally salient facial expressions is widely believed to depend on a subcortical visual circuit from the superior colliculus through the thalamus to the amygdala, but the specific processing steps remain unclear. Here we identified two largely nonoverlapping groups of neurons in the superior colliculus significantly modulated by facial expressions. One group exhibited a short-latency preference for threat and the other exhibited a delayed preference for fear.

## Main Text

Faces are a valuable source of information for primates. Facial expressions are especially important because they provide visual cues about potential danger, social intent, and emotionally salient events. Extracting this information is widely believed to depend on a major subcortical visual circuit through the superior colliculus (SC), pulvinar, and amygdala for quickly detecting faces and identifying threatening and emotionally salient facial expressions^1–3^. This fast and coarse subcortical operation complements the slower but more refined cortical mechanisms for face processing^4–6^ and may play a crucial role in scaffolding the development of the forebrain social networks^7,8^ that are implicated in autism and related disorders^9^. Multiple lines of evidence have established the central role of the amygdala in processing emotionally salient facial expressions^10–12^ but unlike our detailed understanding of the processing steps in visual cortical areas^13–15^, little is known about how these selective responses are constructed. A primary source for facial expression responses in the amygdala is believed to be the SC in the midbrain^12,16^. In fact, visual neurons in the SC themselves exhibit a short-latency preference for faces and face-like stimuli^17,18^. However, it remains unclear whether the encoding of facial expressions is already present at this early midbrain stage or gets added at later processing steps in the pulvinar and amygdala.

To address this gap, we measured visual responses in the SC using images of faces and other object categories, including facial-expression stimuli previously shown to elicit expression-selective activity in amygdala^19^. We recorded the spiking activity of visually responsive neurons in the superficial and intermediate layers of the SC in two macaques while they viewed face and object stimuli, 6° in diameter, centered at either parafoveal (2-6 degrees, Fig. 1A) or foveal (within 2 degrees) locations in the visual field. The image set included four facial expressions (threat, fear, pleasing and neutral), and two nonface categories (hands and human-made objects). Low-level visual features were purposefully varied but matched in distribution across categories including contrast, object size and spatial frequency power in three bands (Fig. S1). As in our previously published work, this strategy allowed us to test whether SC neurons exhibited any systematic preference for faces and facial expressions (or other object categories), independent of low-level visual factors that make the stimuli more or less salient^17^.

**Fig. 1.**
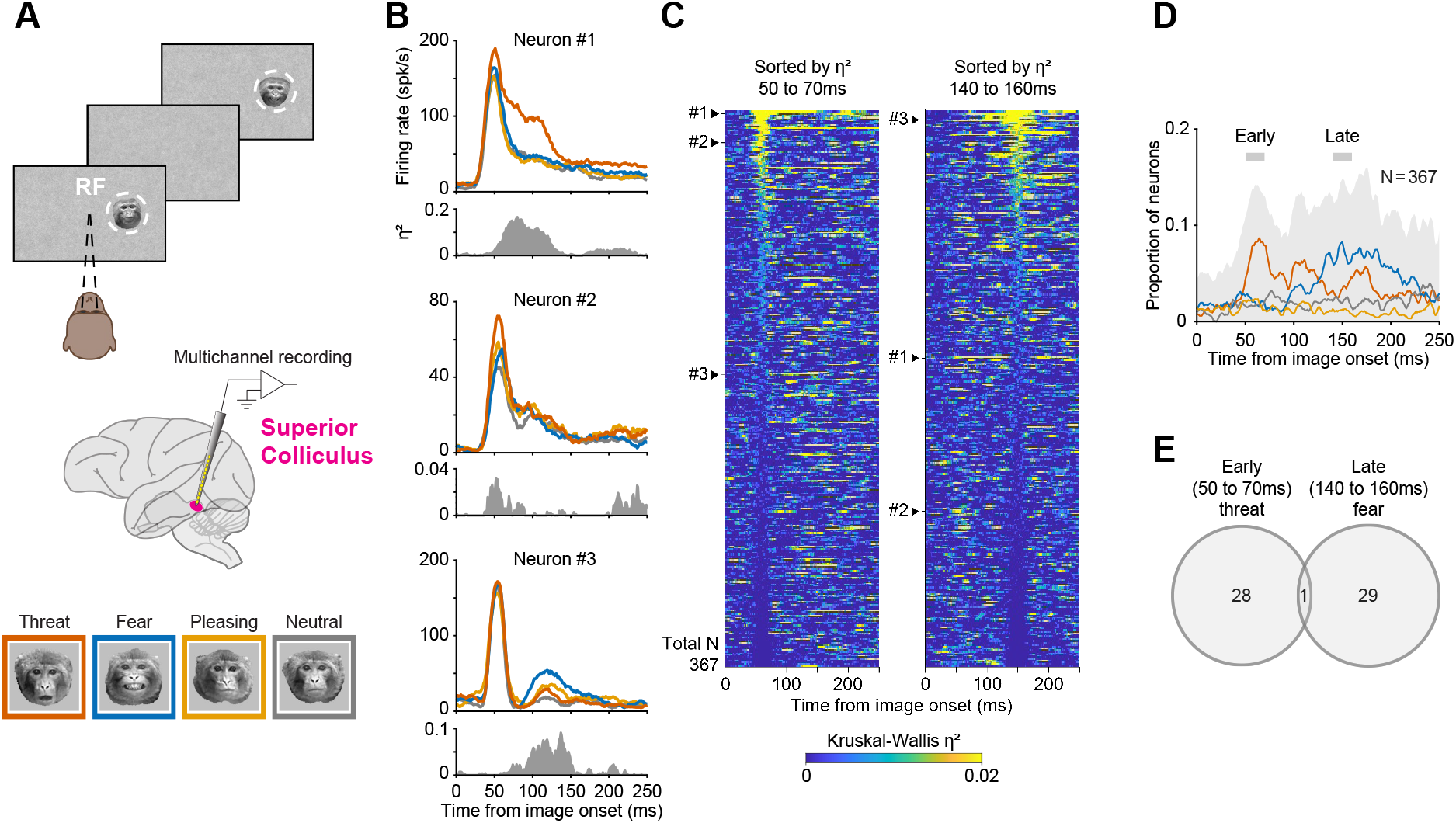
SC neurons are modulated by facial expression during early and late response phases. (A) Experimental paradigm and recording approach. Monkeys maintained fixation while visual stimuli were sequentially presented within the SC response field (RF). Spiking activity was recorded from visually responsive neurons in the superficial and intermediate layers of the SC using multichannel probes. The stimulus set included four facial-expression categories (threat, fear, pleasing, and neutral; 16 images per expression) and two nonface categories (hands and human-made objects). Low-level visual features, including contrast, object size, and spatial frequency power, were matched across categories (Fig. S1). (B) Three example SC neurons showing facial expression-dependent visual responses. For each neuron, trial-averaged firing rates are aligned to image onset and separated by facial-expression category. Below each, gray-filled traces show facial-expression modulation over time, quantified by Kruskal-Wallis η² (eta squared) across the four facial expressions. (C) Expression modulation across all visually responsive SC neurons (n = 367). Heatmaps show η² across time for individual neurons, sorted by mean η² in either the early window (50-70 ms; left) or late window (140-160 ms; right). Neurons #1, #2, #3 are indicated. (D) Time course of the proportion of neurons showing significant facial-expression modulation. Gray shaded area indicates the overall proportion of expression-modulated neurons (preference for any facial expression). Individual colored curves denote the proportion of neurons displaying a preference for a specific facial expression (see legend in A). Early and late windows are indicated by the gray shaded bars. (E) Overlap between neurons with early threat preference and neurons with late fear preference.

We first asked whether individual SC neurons were modulated by facial expressions. Example neurons showed expression-dependent responses during different phases of the visual response (Fig. 1B). Some neurons (neuron #1 and #2) were modulated by facial expression shortly after image onset and responded most strongly to faces expressing threat, whereas others (e.g., neuron #3) were modulated later and exhibited a preference for faces expressing fear. For each neuron, we documented the time course of facial expression modulation using the Kruskal-Wallis effect size, η², across the four facial expressions, providing a statistical measure of the preferences for facial expressions over time (Fig. 1B).

Next, we computed the η² time course for all 367 visually responsive neurons (Fig. 1C). Sorting neurons by η² in an early window (50 to 70 ms) or a late window (140 to 160 ms) produced very different rank orderings of our neurons, suggesting that neurons with strong early modulation generally did not show strong late modulation, and vice versa. We assessed this possibility by quantifying the prevalence and preference of expression-modulated neurons over time (Fig. 1D). During the early phase, centered at 60 ms after image onset, a subset of SC neurons (49 of 367) exhibited a significant effect of facial expression (Kruskal-Wallis H test, p < 0.05, gray shading in Fig. 1D), and these neurons were dominated by a preference for threat faces (29 of 49). During the late phase, centered at 150 ms after image onset, another subset of neurons (58 of 367) exhibited a significant effect of facial expression, but the dominant preference changed to fearful expressions (30 of 58). This early-threat/late-fear pattern was preserved when the analyses were repeated on residual responses after regressing out low-level image features (Fig. S2). The same early-threat/late-fear pattern was also observed when we presented the image set previously used to elicit expression-selective activity in the macaque amygdala, in its original, full color format (Fig. S3)^19^.

We further verified that facial expression preferences were not due to microsaccades, which are known to influence SC activity (Fig. S4). Direct comparison of these effects on a neuron-by-neuron basis revealed minimal overlap between the early threat-preferring and late fear-preferring neurons (Venn diagram in Fig. 1E). Together, these results demonstrate that modulation by facial expression is indeed present in the visual responses of some SC neurons. Moreover, this modulation has a distinctive temporal structure across the population, with early effects of threat and later effects of fear conveyed by largely distinct subsets of SC neurons.

We next investigated how accurately facial expressions could be read out based on SC activity. Although the single-neuron analyses identified a fair number of expression-modulated neurons, they did not establish whether population activity contained enough information to discriminate between different facial expressions. To test this, we trained linear classifiers using pseudopopulation responses computed in 40-ms bins from all visually responsive SC neurons and evaluated their cross-validated accuracy for each pairwise classification of facial expressions (Fig. 2A). The performance of these pairwise classifiers was markedly different in the early and late phases of the visual response. In the early window, 40 to 80 ms after image onset, the strongest pairwise performance involved discriminating threat faces from nonnegative expressions, including pleasing and neutral faces (Fig. 2B, left). In contrast, in the late window, 130 to 170 ms after image onset, decoding performance was best for discriminating fear faces from nonnegative expressions (Fig. 2B, right).

**Fig. 2.**
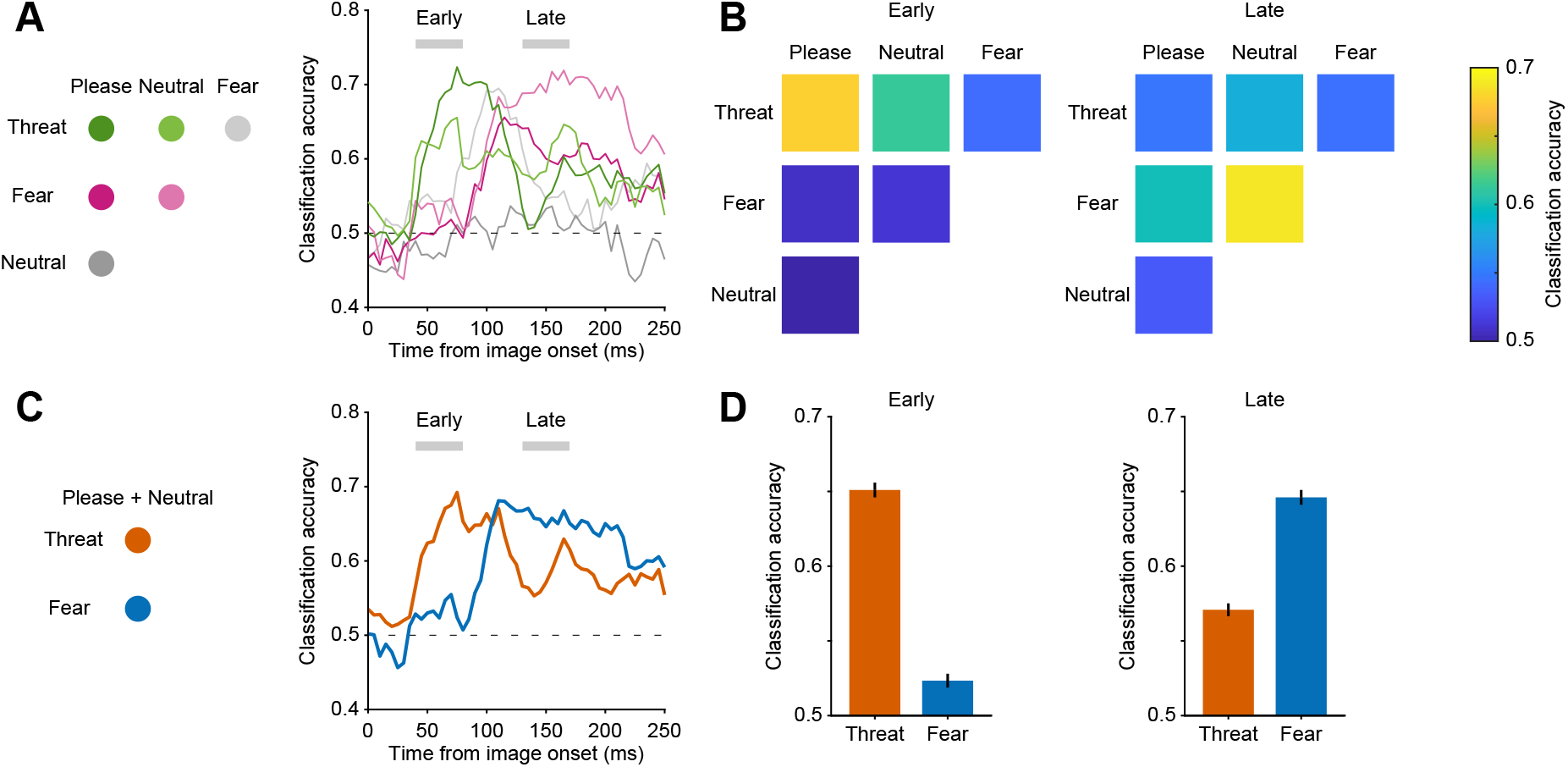
SC population decoding reveals early threat and late fear signals. (A) Pairwise population decoding of facial-expression category from SC pseudopopulation activity. Linear classifiers were trained on responses from all visually responsive neurons (n = 367) and tested using cross-validation. Classification accuracy is shown for each pairwise comparison among threat, fear, pleasing, and neutral faces. Horizontal dashed line indicates chance performance; gray shaded bars indicate the early (40-80ms) and late (130-170ms) decoding windows for constructing confusion matrices in panel B. (B) Confusion matrix summary of pairwise decoding accuracy in the early and late windows. In the early window, accuracy was highest for decoding threat faces from pleasing faces. In the late window, accuracy was highest for decoding fear faces from neutral faces. (C) Time-resolved binary decoding of threat/fear faces versus the nonnegative expressions (pleasing and neutral faces). (D) Comparison of decoding accuracy for threat and fear classifiers in the early and late windows.

We next used targeted binary classifiers to directly decode either threat faces or fear faces against the same nonnegative expressions (Fig. 2C). Threat-related information was strongest in the early response window, whereas fear-related information became stronger later in the response, matching the temporal structure observed at the single-neuron level (Fig. 2D). Thus, information about both threat- and fear-related facial expressions could be read out from SC population activity, with threat information emerging earlier and fear information becoming more prominent later in the visual response.

Finally, we considered whether facial-expression signals differed between SC neurons with foveal and parafoveal receptive fields, because neurons representing different visual eccentricities might have different functional roles in face processing^20^. SC neurons with eccentric receptive fields are well positioned to detect faces outside the center of gaze and guide orienting movements toward them. Foveal SC neurons, in contrast, respond once a face is directly fixated, placing the stimulus in a social context potentially shaped by individual differences in dominance and social experience. We therefore compared expression-related activity and population decoding performance for foveal and parafoveal SC neurons (Fig. 3A). At the single-neuron level, the proportion of expression-modulated neurons differed between foveal and parafoveal neurons, with the largest differences occurring for threat- and fear-preferring neurons (Fig. 3B). Population decoding revealed that threat-related information was present in both foveal and parafoveal neurons during the early response window, but only the parafoveal population maintained strong threat-related information into the middle and late response windows (Fig. 3C). Fear-related information followed a different temporal profile: it was weak in the early response window, became stronger later, and was most prominent in the parafoveal population during the late response window (Fig. 3D). The difference in decoding accuracy, computed as foveal minus parafoveal accuracy, emphasized these distinct temporal profiles (lower plots in Fig. 3C,D). We also examined these effects in each animal separately (Fig. S5). The parafoveal pattern was evident in both monkeys, with early threat-related modulation, late fear-related modulation, and sustained threat-related information over time. Foveal expression-related signals were more animal dependent: the older monkey (#1; 16-year-old) showed little threat-related information but strong fear-related information for foveal presentations, whereas the younger monkey (#2; 8-year-old) showed stronger foveal threat-related information. Nevertheless, the most robust pattern across animals was found in parafoveal SC, where threat-related information emerged early and persisted, and fear-related information emerged later.

**Fig. 3.**
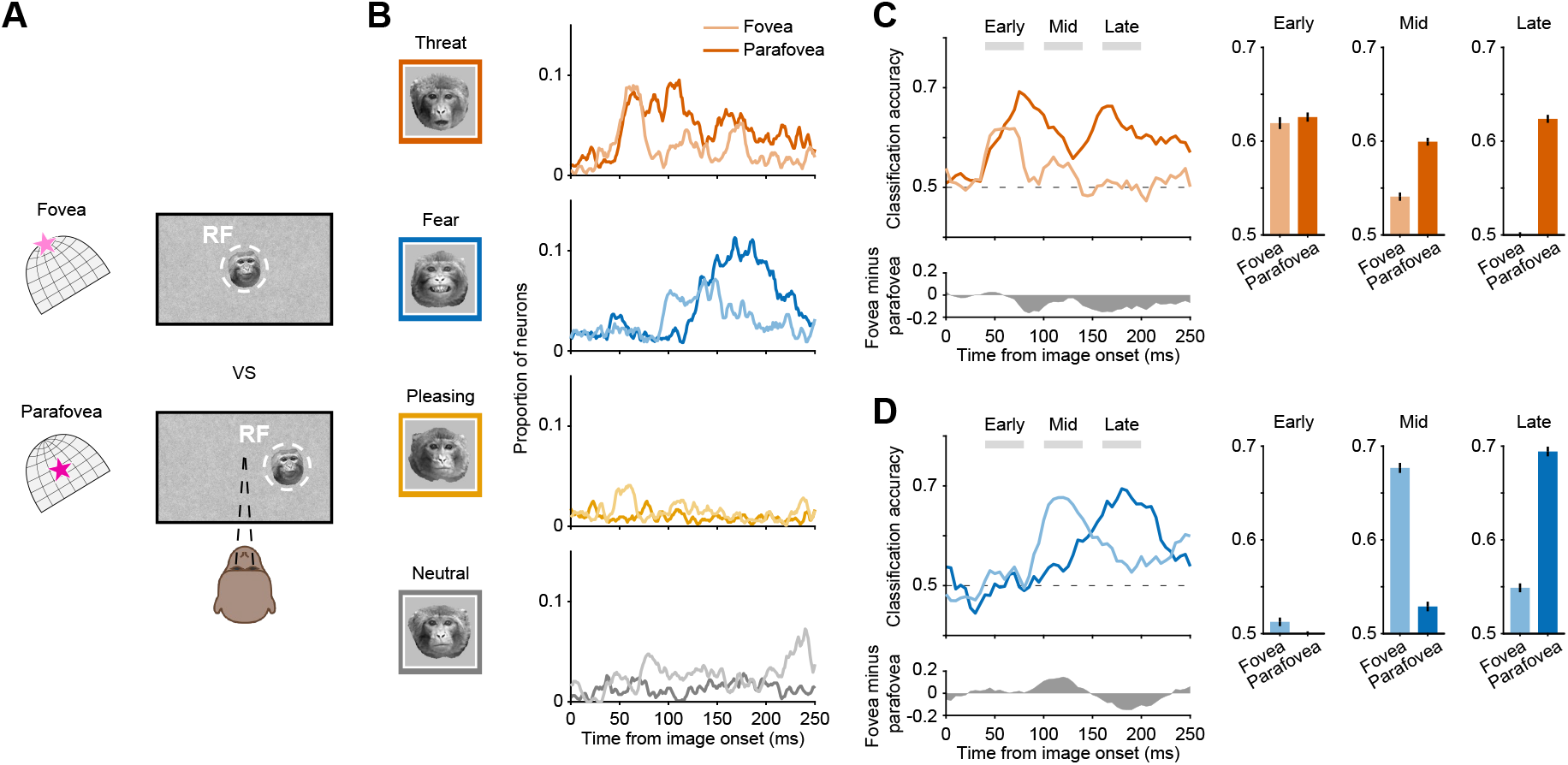
Threat and fear signals show distinct temporal profiles between foveal and parafoveal SC neurons. (A) Comparison of foveal and parafoveal SC responses. Recordings were made from either rostral or caudal SC (magenta star on SC schematic). Visual stimuli were presented either at foveal RFs, defined as locations within 2 degrees of fixation, or at parafoveal RFs, defined as locations 2-6 degrees from fixation. (B) Time course of single-neuron expression preference for foveal and parafoveal SC neurons. Traces indicate the proportion of neurons preferring each facial-expression category among neurons with significant expression modulation. Lighter traces indicate foveal responses, and saturated traces indicate parafoveal responses. (C) Population decoding of threat faces versus nonnegative expressions for foveal and parafoveal neurons. Gray shaded bars indicate early, middle, and late decoding windows. Bar plots summarize decoding accuracy in each window. The lower trace shows foveal minus parafoveal decoding accuracy; positive values indicate stronger foveal decoding and negative values indicate stronger parafoveal decoding. (D) Population decoding of fear faces versus nonnegative expressions for foveal and parafoveal presentations, plotted in the same format as panel C.

These results provide a key piece of information about the subcortical circuits for rapid evaluation of emotionally salient facial expressions. Because emotional face processing depends strongly on low spatial frequencies that are also preferred by neurons in the superior colliculus^21^, it has been speculated that a rudimentary form of facial-expression preference might already be computed at this early stage in the midbrain^22^. Our results provide, to our knowledge, the first single-neuron evidence supporting this idea in the primate SC. Moreover, our results show that this expression-related information has a distinctive temporal and spatial structure, with an earlier threat-related component and a later fear-related component, represented by largely distinct subsets of neurons. This dissociation suggests that threat and fear are processed differently in these subcortical circuits. The threat-related component was especially robust and sustained in parafoveal SC, where it is well positioned to detect potential danger outside the center of gaze and could drive rapid behavioral responses to avoid harm. Fear-related activity, by contrast, emerges later, consistent with an expression-related signal that may involve interactions between SC and broader face-processing circuits. Identifying how these signals contribute to the properties in other subcortical and cortical areas will be important for understanding the development and function of visual pathways for social perception across neurotypical and neurodivergent populations^5,7,8^.

## Acknowledgements

We thank Daniel Yochelson, Nick Nichols, Denise Parker, and Hayden Warnock for technical support. We are grateful to members of the Krauzlis lab for helpful discussions and feedback. We also thank Katalin Gothard for generously providing the macaque facial-expression image set. This work is supported by the National Eye Institute Intramural Research Program at the National Institutes of Health ZIA EY000511. The contributions of the NIH authors were made as part of their official duties as NIH federal employees, are in compliance with agency policy requirements, and are considered Works of the United States Government. However, the findings and conclusions presented in this paper are those of the author(s) and do not necessarily reflect the views of the NIH or the U.S. Department of Health and Human Services.

## Supplementary Figures

**Fig. S1.**
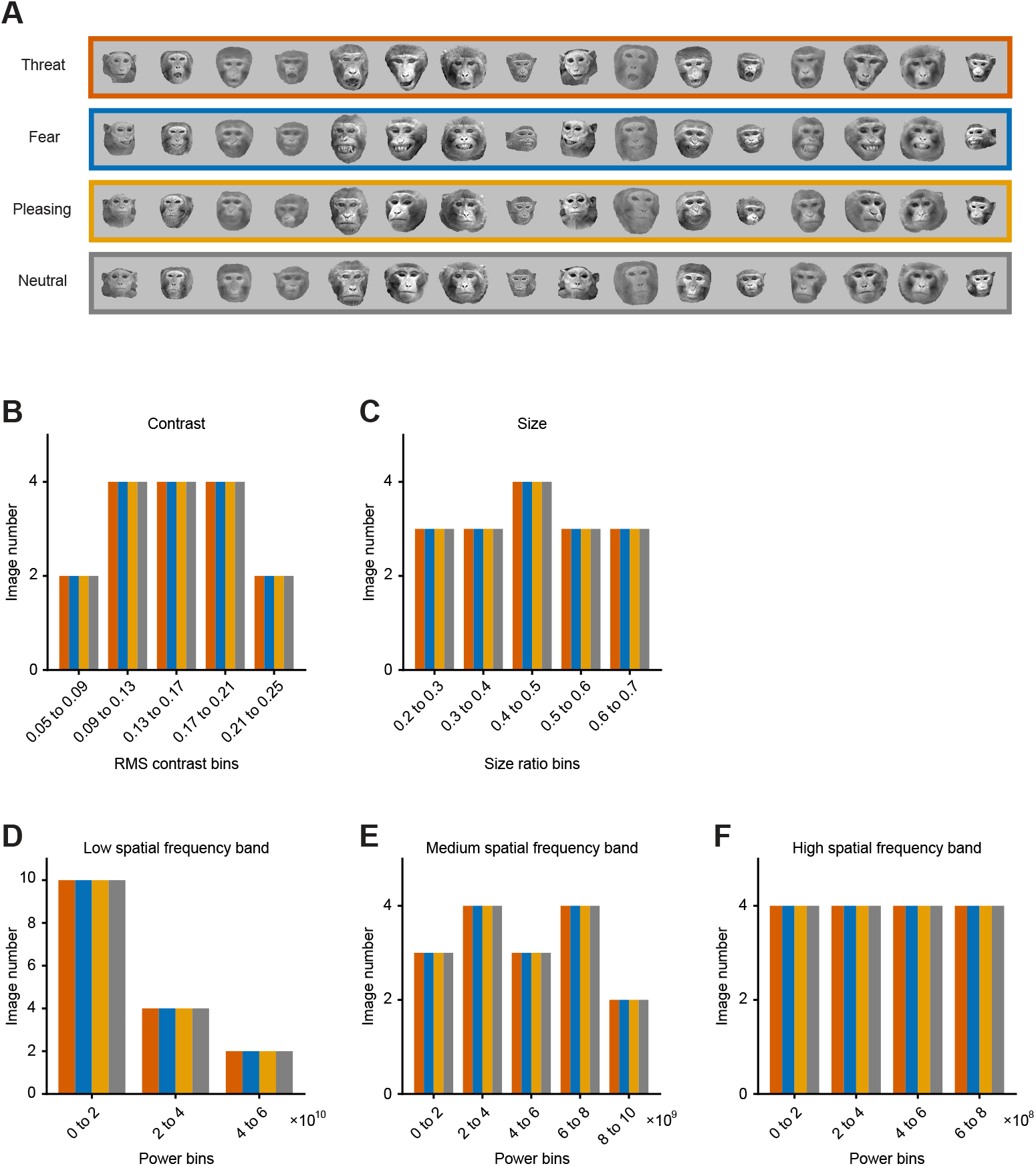
Distributions of low-level visual features were matched across object categories. (A) The facial expression image set consisted of 64 images, 16 each from four facial expression categories (threat, fear, pleasing and neutral). Across the four facial expressions, the distribution of five low-level features was matched. The features are: (B) RMS contrast; (C) Size ratio (number of object pixels divided by number of pixels in the 6 deg aperture); (D) Spectral power in a low spatial frequency (SF) band (0.1667 to 0.6525 cycles/degree); (E) Spectral power in an intermediate SF band (0.6525 to 2.5544 cycles/degree); (F) Spectral power in a high SF band (2.5544 to 10 cycles/degree). Distribution matching was confirmed with the Kolmogorov–Smirnov test (p > 0.05, for each of the possible pairwise comparisons of object categories for each low-level feature). Additionally, mean luminance of all object images was matched to the mean luminance of the background (32.6cd/m^2^).

**Fig. S2.**
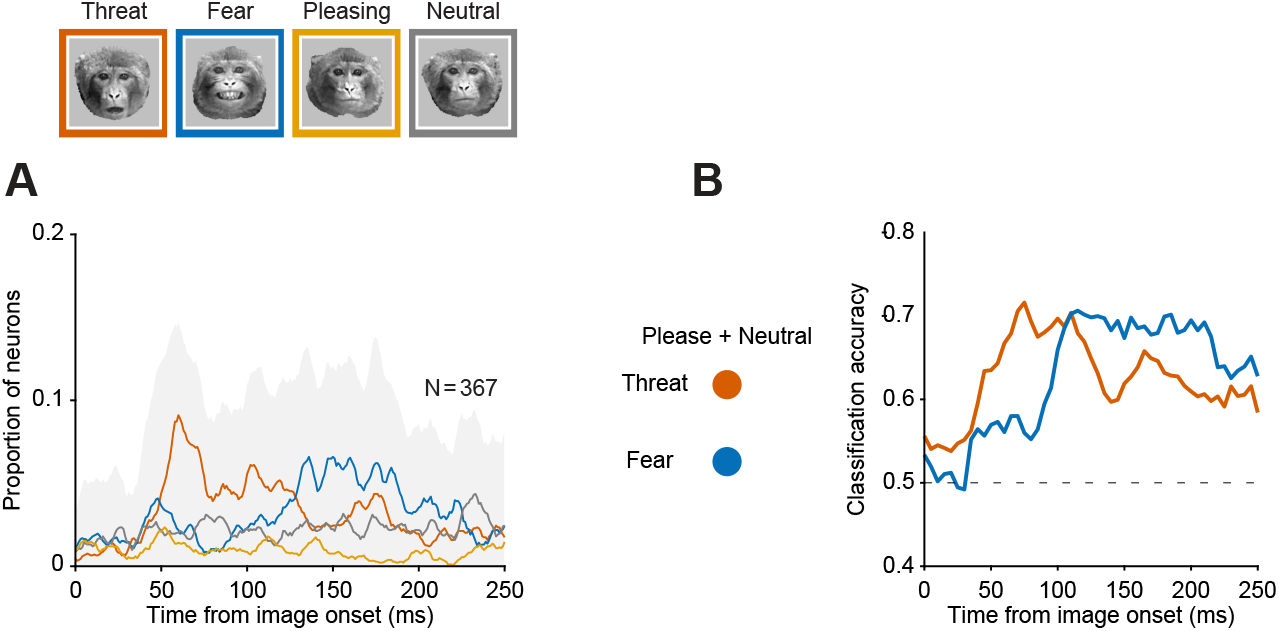
Facial expression preferences in SC persists after regressing out low-level visual features. (A) Proportion of SC neurons exhibiting facial-expression preferences over time after removing the contribution of low-level visual features. For each neuron and time bin, responses were modeled using a multilinear regression with five image features as regressors: RMS contrast, size ratio, and power in low, medium, and high spatial-frequency bands. Facial-expression preference was then assessed on the residual responses using a Kruskal–Wallis H test with facial expression as the group variable (p < 0.05). Gray shaded area indicates the overall proportion of expression-modulated neurons (preference for any facial expression). Individual colored curves denote the proportion of neurons displaying a preference for a specific facial expression. (B) Cross-validated decoding accuracy computed from the residual responses. Linear classifiers were trained to discriminate threat faces from nonnegative expressions and fear faces from nonnegative expressions, where nonnegative expressions included pleasing and neutral faces.

**Fig. S3.**
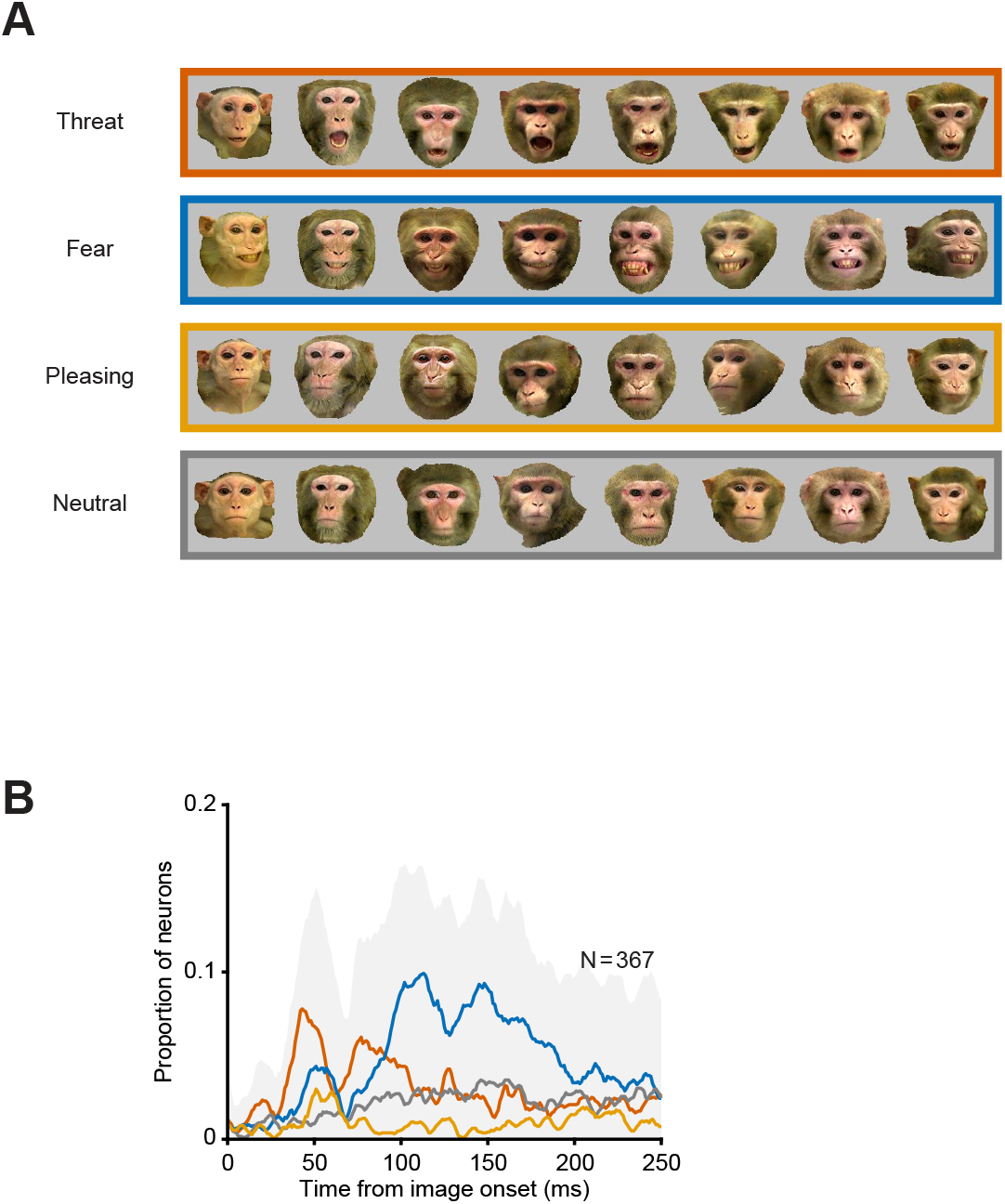
Early threat- and late fear-related pattern was preserved for original color images. (A) Original color facial-expression image set previously shown to elicit expression-selective neural activity in the macaque amygdala. (B) Time course of the proportion of neurons showing significant color-facial-expression preference. Gray shaded area indicates the overall proportion of expression-modulated neurons (preference for any facial expression). Individual colored curves denote the proportion of neurons displaying a preference for a specific facial expression.

**Fig. S4.**
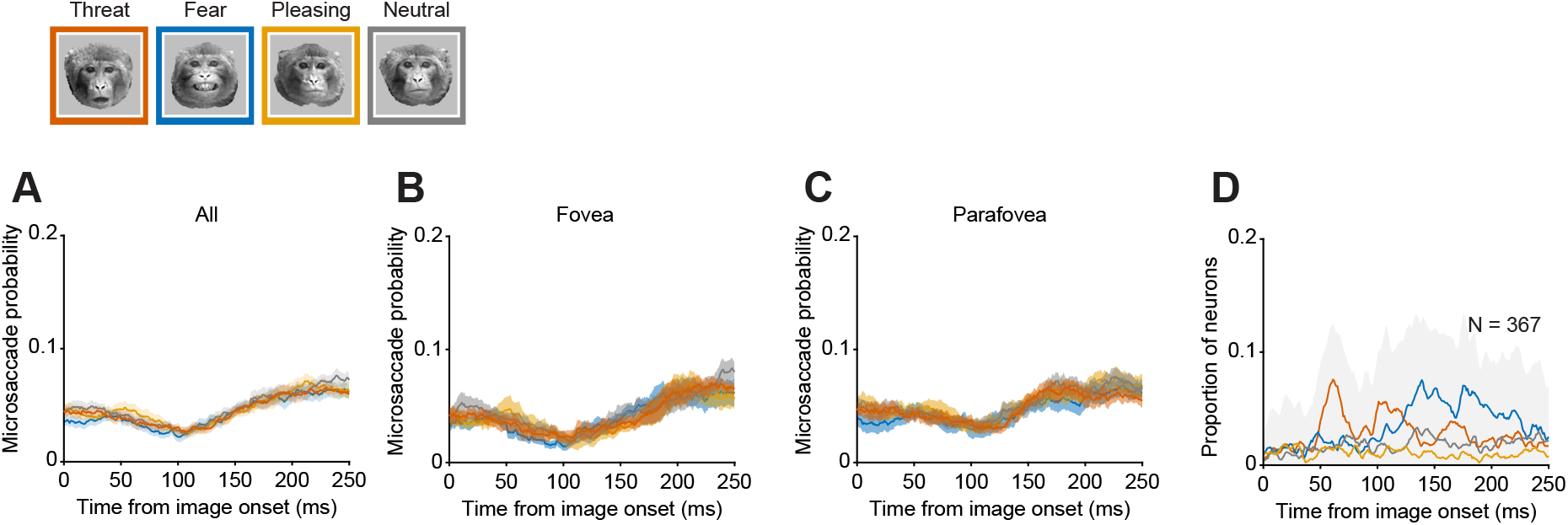
Facial expression preferences in SC cannot be explained by the occurrence of microsaccades. Probability of microsaccades over time in 20ms non-overlapping bins aligned to image onset for the four facial expressions, shown for all trials (A), foveal conditions (B), and parafoveal conditions (C). There was no significant difference in microsaccade probability across facial expressions in any time bin (Kruskal–Wallis H test with facial expression as the group variable, p > 0.05). (D) Proportion of SC neurons exhibiting facial-expression preferences (Kruskal–Wallis H test with facial expression as the group variable, p < 0.05) after excluding trials containing microsaccades from 0 to 250 ms after image onset. Gray shaded area indicates the overall proportion of expression-modulated neurons (preference for any facial expression). Individual colored curves denote the proportion of neurons displaying a preference for a specific facial expression.

**Fig. S5.**
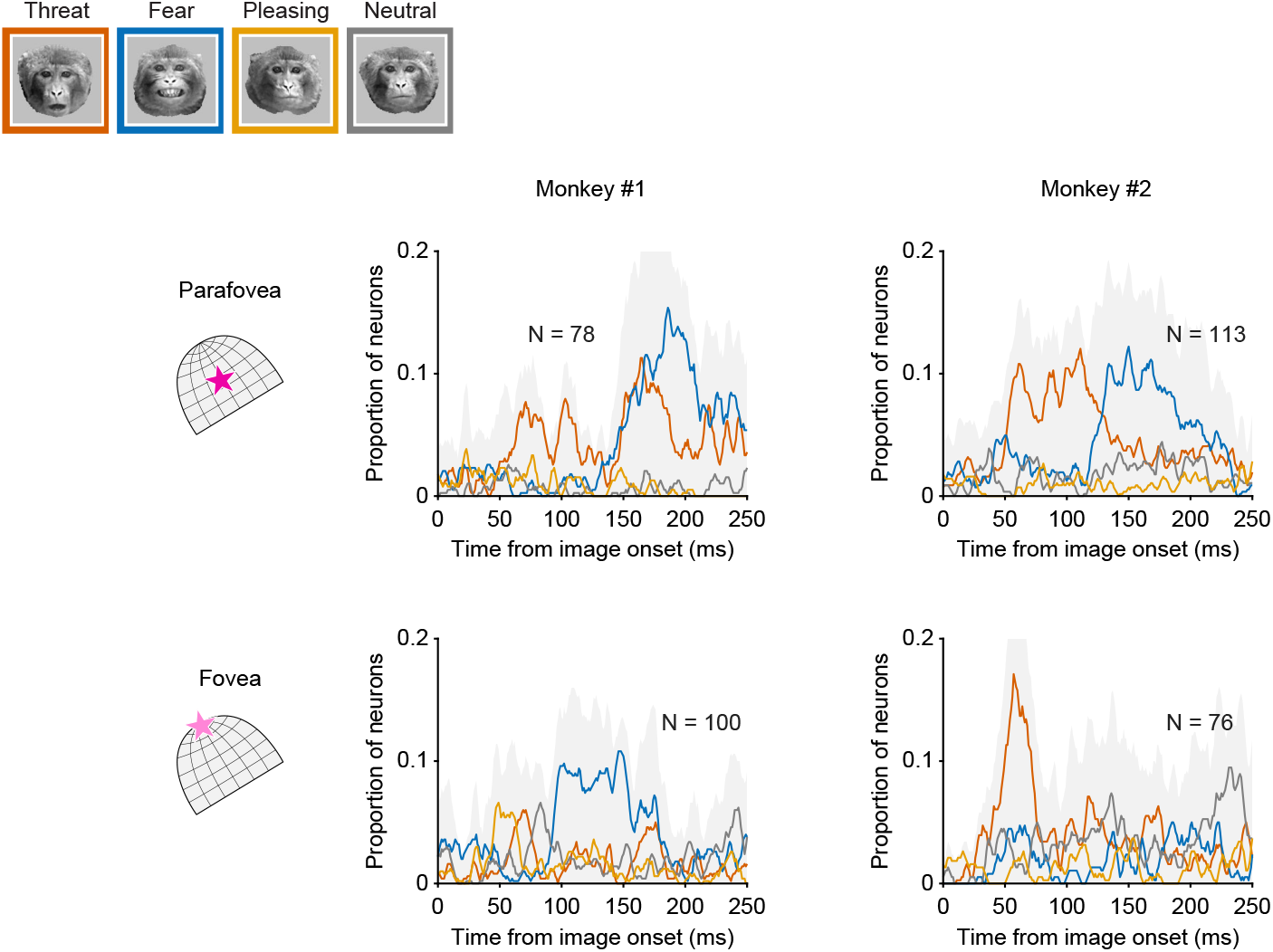
Expression-related signals in individual animals. Time courses show the proportion of neurons with significant facial-expression modulation, plotted separately for monkey 1 and monkey 2. Gray shading indicates the overall proportion of expression-modulated neurons, defined as neurons with a significant effect of facial expression regardless of preferred expression. Colored curves indicate the proportion of neurons preferring each facial expression. Top row shows parafoveal visually responsive neurons and the bottom row shows foveal neurons. Across animals, the most robust shared pattern was found in parafoveal SC, where threat-related modulation emerged early and persisted, whereas fear-related modulation emerged later. Foveal expression-related signals showed individual-specific differences in the relative strength of threat- and fear-related information.

## Methods

### Experimental model and study participant details

Two adult male rhesus macaques (Macaca mulatta) participated in the study: monkey #1 was 16 years old and weighed 12 kg, and monkey #2 was 8 years old and weighed 8 kg. All experimental procedures were approved by the National Eye Institute Animal Care and Use Committee and were conducted in accordance with the United States Public Health Service Policy on Humane Care and Use of Laboratory Animals. Each animal had previously been implanted with a plastic headpost and recording chamber to permit electrophysiological access to the superior colliculus (SC).

### Experimental apparatus

Monkeys were seated in a custom primate chair (Crist Instrument Co., Hagerstown, MD) and head-fixed in a darkened booth facing a VIEWPixx display (VPixx Technologies; 100 Hz refresh rate). Display latency was measured with a photodiode, and all event times were corrected accordingly. Experiments were controlled using a modified version of the PLDAPS framework^1^. Horizontal and vertical eye position were recorded at 1,000 Hz using an EyeLink infrared eye-tracking system.

### Saccade tasks

Monkeys performed standard visually guided saccade tasks to map neuronal receptive fields (RFs). In the visually guided saccade task, monkeys acquired central fixation, and a peripheral target appeared 0.75–1 s later and remained visible. After a delay of 1–2 s, the fixation point disappeared, providing the go cue to saccade to the target. Target locations were varied across trials to estimate the location and extent of each neuron’s RF. A memory-guided version of the task was used to classify neurons as either visual, visual-movement, or movement neurons based on their responses to target appearance and saccade execution. In the memory-guided condition, the target was reillumined following entrance into the target window for a duration of 0.2 s. Additional task and classification details have been described previously^2^.

### Object viewing task

Following RF mapping, the monkeys performed an object-viewing task while maintaining central fixation. Trials began when the monkey pressed a joystick, triggering the appearance of a 0.25° white fixation square (48 cd/m²) on a uniform gray background (32.6 cd/m²). Eye position was required to remain within an invisible 1.5° square fixation window. After fixation was acquired, the gray background was replaced by a pink-noise texture of the same mean luminance (root-mean-square contrast, 4.38%). Following a 500ms delay, 3 to 5 visual objects (selected pseudo-randomly on every trial) were sequentially presented at a prescribed location (0-6° eccentricity depending on the receptive field locations of the recorded neurons). Each stimulus was displayed for 400ms and was followed by a 400ms interstimulus interval. The pink-noise background was refreshed at each stimulus onset and offset. Stimuli were presented within a 6° diameter circular aperture regardless of visual-field location. The monkey received a liquid reward after maintaining fixation throughout the complete stimulus sequence. Only successfully completed fixation trials were included in the analyses. Overall, monkeys completed 226 ± 30 (mean ± sd) trials across sessions, and each object was presented 7 ± 1 times. Only data from successfully completed trials were analyzed.

### Image set

The image set included grayscale images of four macaque facial expressions: threat (open-mouth threat), fear (fear grin), pleasing (lipsmacking), and neutral. The original image set comprised 32 images from eight individual monkeys, with one image of each facial expression available for each monkey identity. To increase the number of exemplars while preserving the expression and identity structure of the stimulus set, each image was also horizontally mirrored, yielding 64 face images in total (8 monkey identities, 8 images per identity, netting 16 images per facial expression category). The facial expression stimuli were grayscale versions of images previously used to demonstrate expression-related neuronal activity in the macaque amygdala^3^. Images of hands and human-made objects were also presented during the same recording sessions, with 30 images in each nonface category.

As in our previous study^4^, low-level image properties were matched in distribution across stimulus categories. Images were processed using the SHINE toolbox^5^ and custom MATLAB code to control mean luminance, RMS contrast, image size, and power in low, medium and high spatial frequency bands across categories (for precise values ranges see Fig. S1). The distributions of these features were matched across the four facial-expression categories and across the two nonface categories. In addition, the four facial-expression images from each monkey identity were matched in their low-level visual features, minimizing systematic covariation between facial expression and basic visual properties both across the image set and within identity.

Matching the low-level image features across image categories already allowed us to interpret any differential response in SC neurons as driven by the image category, and not by image features. However, some variation in low-level features still exists across images. To account for their potential effect on SC neurons’ responses we constructed a multiple linear regression model (as in^4^). For each neuron and time bin, responses were modeled using a multilinear regression with five image features as regressors: RMS contrast, size ratio, and power in low, medium, and high spatial-frequency bands. We then assessed the facial-expression preference on the residual responses using a Kruskal–Wallis H test with facial expression as the group variable (p < 0.05), and also performed population decoding analyses on the residuals (Fig. S2).

### Electrophysiology recordings

Electrophysiological signals were acquired using an OmniPlex-D system (Plexon Inc.). Neuronal activity was recorded from the superficial and intermediate layers of the SC using 32-channel Plexon V-probes with 50-µm interchannel spacing. Probes were advanced with a motorized Microdrive (NAN Instruments) to depths containing visually responsive and visual-movement neurons. Probe position was adjusted to maximize neuronal yield and was then allowed to stabilize for approximately 1 h before data collection. Because the probe entered the SC obliquely, receptive fields recorded simultaneously generally overlapped; the stimulus location was selected to maximize overlap between the 6° aperture and the recorded receptive fields. Overall, recordings took place over 29 sessions in the two monkeys.

### Electrophysiology analysis

Continuous spike-channel data were sorted offline using Kilosort2^6^ and in-house tools (https://github.com/ElKatz/kilo2Tools) and were manually curated in Phy2. Units were retained only when their waveform shapes and inter spike-interval distributions were consistent with well-isolated action potentials. Neurons with a low (<1.8) signal-to-noise ratio or low average trial firing rate (<1 spikes/s) were excluded. SC neurons were then classified according to their activity during a memory-guided saccade task using standard criteria^2,7^, and only visually responsive neurons were included in the analyses. Overall, 624 neurons were recorded (357 from Monkey #1, 267 from Monkey #2) and 257 of these did not meet our inclusion criteria, netting 367 neurons for the analysis reported here (178 from Monkey #1, 189 from Monkey #2). Data were combined across monkeys for the main analyses to increase statistical power. Individual-monkey analyses largely supported the key findings and are presented in Fig. S5.

For visualization of time-resolved mean firing rates relative to key events in the task (Fig. 1B), spike counts were computed in overlapping 20ms bins advanced in 1ms steps and aligned to stimulus onset.

Facial expression modulation was quantified for each neuron over time (Fig. 1D, S2A, S3B, S4D, S5) using a one-way non-parametric ANOVA (Kruskal-Wallis H test) with facial-expression category (threat, fear, pleasing, or neutral) as the main factor, on spike counts for all facial expression images in overlapping 20ms bins sliding in 1ms steps aligned to stimulus onset. A neuron was considered facial expression modulated in time bins for which p < 0.05. The facial expression which induced the largest neuronal responses was defined as the preferred facial expression of the neurons. The magnitude of expression modulation was summarized with the Kruskal-Wallis η² effect size (Fig. 1B and 1C).

### Microsaccade detection

Microsaccades were detected during the object viewing task using a 2D-velocity-based algorithm (relative velocity threshold = 6 and minimum saccade duration = 6ms) developed by^8,9^.

### Binary decoders for facial expression classification

We used cross-validated binary linear classifiers to determine whether population responses of SC neurons contained information that could discriminate facial expressions. We performed all six pairwise comparisons of the four facial expression categories (Fig. 2A, legend). We additionally tested whether threat or fear expressions could be discriminated from the combined set of nonnegative expressions, comprising pleasing and neutral faces (Fig. 2C, legend).

For each classification repeat, images from each of the two classes under consideration were randomly divided into nonoverlapping training and testing sets, with 70% of the images assigned to training and the remaining 30% assigned to testing. Classifier performance was therefore evaluated using image identities that had not been included during training, requiring the classifier to generalize across exemplars of the same facial-expression category.

Neuronal responses were measured as spike counts within a 40-ms analysis window. Because neurons were recorded across different sessions rather than simultaneously, population response vectors were constructed as stimulus-locked pseudotrials. For each pseudotrial, one image was first selected from the relevant training or testing image set. For each neuron, one response to that same image was then sampled randomly, with replacement, from the available repetitions. For each classification repeat, we generated 1,000 training pseudotrials and 1,000 testing pseudotrials for each class, yielding 2,000 training and 2,000 testing population response vectors. A support vector machine with a linear kernel was trained on the training pseudotrials and evaluated on the testing pseudotrials. Accuracy was calculated as the proportion of correctly classified testing pseudotrials. The complete procedure, including image-level training–testing partitioning, pseudotrial generation, model fitting, and testing, was repeated 100 times. We report the mean classification accuracy across repeats, with the standard error of the mean.

For the principal decoding analyses shown in Figure 2, classifiers were constructed using all visually responsive SC neurons included in the analysis. For the comparison between foveal and parafoveal SC populations shown in Figure 3, the two populations contained different numbers of neurons. We therefore matched population size by randomly subsampling neurons from the larger population to equal the number available in the smaller population. A new matched neuronal sample was selected for each classification repeat, and the same selected neurons were used to construct all training and testing pseudotrials within that repeat.

To characterize the time-varying accuracy of facial expression classification, independent classifiers were trained and tested in overlapping 40-ms windows advanced in 5-ms steps.

